# Reduced social function in experimentally evolved *Dictyostelium discoideum* implies selection for social conflict in nature

**DOI:** 10.1101/2023.09.02.556041

**Authors:** Tyler J. Larsen, Israt Jahan, Debra A. Brock, Joan E. Strassmann, David C. Queller

## Abstract

Many microbes interact with one another, but the difficulty of directly observing these interactions in nature makes interpreting their adaptive value complicated. The social amoeba *Dictyostelium discoideum* forms aggregates wherein some cells are sacrificed for the benefit of others. Within chimeric aggregates containing multiple unrelated lineages, cheaters can gain an advantage by undercontributing, but the extent to which wild *D. discoideum* has adapted to cheat is not fully clear. In this study, we experimentally evolved *D. discoideum* in an environment where there were no selective pressures to cheat or resist cheating in chimeras. *D. discoideum* lines grown in this environment evolved reduced competitiveness within chimeric aggregates and reduced ability to migrate during the slug stage. By contrast, we did not observe a reduction in cell number, a trait for which selection was not relaxed. The observed loss of traits that our laboratory conditions had made irrelevant suggests that these traits were adaptations driven and maintained by selective pressures *D. discoideum* faces in its natural environment. Our results suggest that *D. discoideum* faces social conflict in nature, and illustrate a general approach that could be applied to searching for social or non-social adaptations in other microbes.

**SIGNIFICANCE STATEMENT:** Microbes interact in diverse and important ways, but the difficulty of directly observing microbes in nature can make it challenging to understand the adaptive significance of these interactions. In this study, we present an experimental evolution approach to infer the selective pressures behind an apparently social trait in the microbe *Dictyostelium discoideum.* We take advantage of the observation that organisms ‘use it or lose it’ – when selective pressures are relaxed, adaptations that evolved in response to those pressures tend to be lost. Our work helps resolve debate over the importance of cheating in *D. discoideum,* and demonstrates a general approach that could be applied to the study of other microbial traits that are difficult to observe in nature.

## INTRODUCTION

Microbes are capable of social behavior that once would have seemed beyond the abilities of such tiny organisms. Despite their small size and simplicity, microbes can cooperate to sense their environment [1–3], hunt prey [4], kill enemies [5], protect friends [6], move over difficult ground [7], collect nutrients [8], and more. Microbial cooperation is worthy of study in its own right for its significant consequences for human health and on the ecological services microbes provide, but also because microbes have proven to be uniquely valuable model organisms for studying major questions about evolution [9–11].

One long-standing question of interest is how cooperation evolves and is maintained despite the threat of exploitation by non-cooperating cheaters [12–16]. Many functions that larger organisms perform privately microbes must perform publicly through the production and secretion of molecules into their environment, which would seem to render them especially vulnerable to the risk of exploitation. This, combined with their short generations and high population sizes, makes social microbes especially well-suited for studying the evolution of cooperation and conflict.

*Dictyostelium discoideum* is an interesting eukaryotic microbe with utility as a model organism for scientists studying development, multicellularity, immunology, and cooperation and conflict [17–19]. While it spends most of its life as a solitary unicellular hunter of bacteria, when starving, *D. discoideum* enters a multicellular life stage **(Figure 1)** [19–21]. Local amoebae aggregate together to form a slug-like multicellular body that can move towards light. Upon reaching a suitable spot, the slug develops into a fruiting body consisting of a ball-shaped sorus of durable spores held aloft by a stalk of dead somatic cells. The spores wait dormant in the sorus to be dispersed – possibly by a passing invertebrate – to a new location with sufficient prey [22, 23].

**FIGURE 1.**
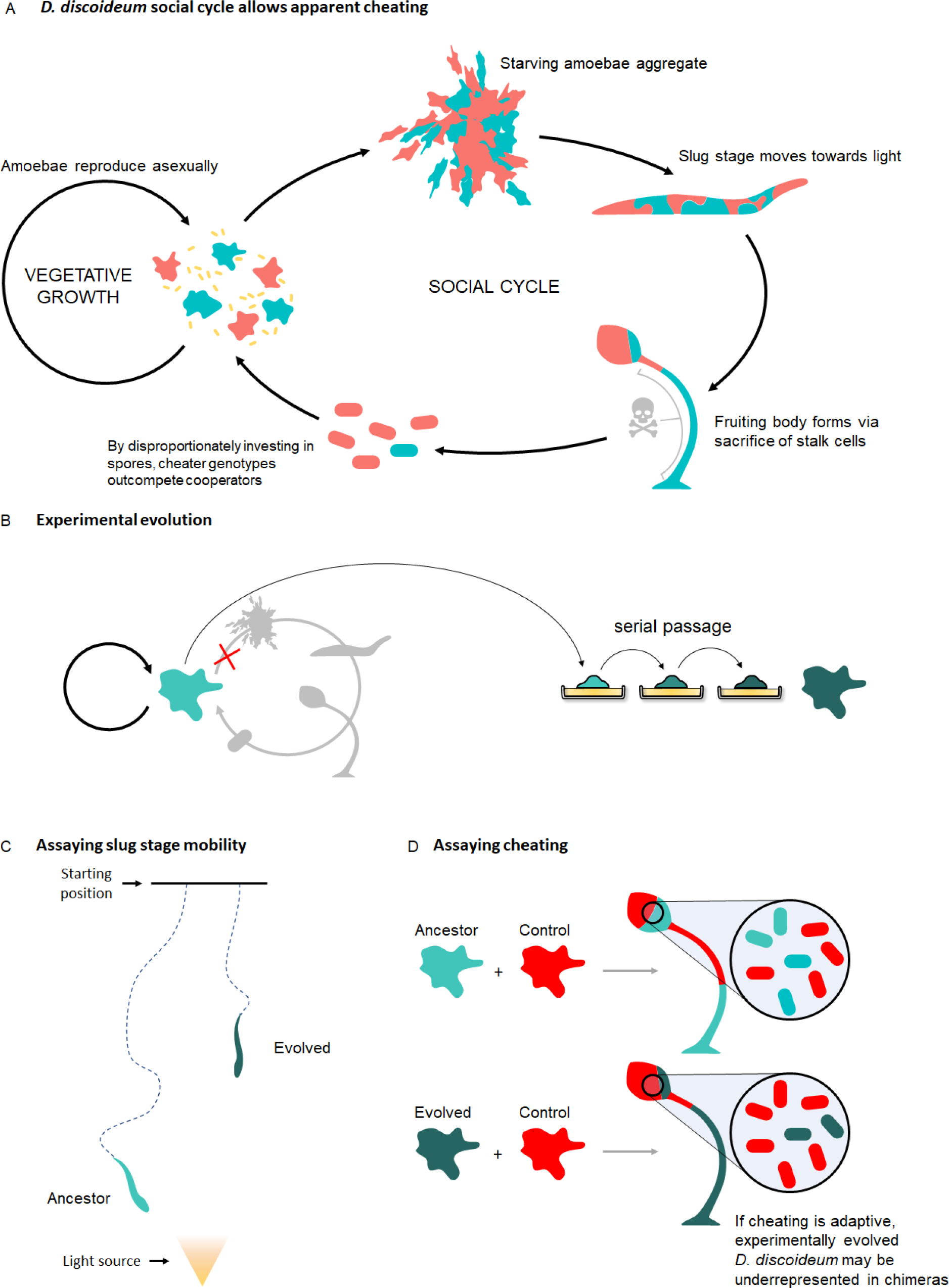
Experiment Overview. **A)** simplified schematic of *D. discoideum’s* social cycle, in which multiple genotypes (red and teal) can potentially aggregate into chimeric multicellular bodies. Most of the time, amoebae are unicellular and grow vegetatively. Upon starvation, cells aggregate and develop into a multicellular slug stage, which moves towards light, and then a sessile fruiting body. Formation of the fruiting body requires the sacrifice of a minority of cells to produce a stalk, and so cheater genotypes (red) can gain an advantage by undercontributing to stalk formation. Cheating appears to provide clear benefits in artificial settings but its relevance to wild *D. discoideum* is debated. **B)** Experimental evolution of *D. discoideum* under conditions which prevent it from entering the social cycle should drive the loss of traits previously maintained by selective pressures related to it. **C)** Slug mobility assays - Experimentally evolved *D. discoideum* (dark green) should travel less far during the slug stage than its ancestor (teal) due to relaxed selection. **D)** Cheating assays - Experimentally evolved *D. discoideum,* when mixed equally into a chimera with an RFP-labelled control strain (red), should be less well represented among the spores of the resulting fruiting body than its ancestor due to relaxed selection.

*Dictyostelium discoideum* aggregates form from local amoebae, which do not need to be closely related [18, 24, 25]. When cells aggregate into chimeras containing multiple unrelated cell lineages, they have the opportunity for conflict that cells within more conventional multicellular organisms – made up of the clonal descendants of a single cell – mostly avoid. In *D. discoideum* this conflict centers on the production of its fruiting body’s characteristic stalk, which requires about 20% of the cells within an aggregate to die to help disperse the remainder. Such a sacrifice would not be remarkable within a clone (picture the staggering majority of human cells that toil and die just to pass on a few gametes), but in a chimeric aggregate, it creates an incentive for competing cell lineages to under-contribute to stalk production to minimize their own losses and exploit more civically-inclined lineages [18, 26, 27].

Multiple studies have observed cheating in *D. discoideum* [18, 28–31]. *D. discoideum* chimeras form readily in the laboratory, and careful observation of the fates of cells within them reveals that not every cell lineage contributes equally to stalk production. Some *D. discoideum* strains appear to be consistently prone to exploiting or prone to being exploited by others [28, 32] such that a simple hierarchy of cheaters and cooperators can be determined. Studies have identified mutations that cause strains to cheat [29–31, 33], and in experimental evolution experiments where *D. discoideum* is allowed to mutate, mix, and fruit, non-fruiting obligate cheaters readily evolve and outperform cooperative strains, even to the point of causing population crashes [34]. Some evidence suggests that *D. discoideum* has mechanisms for maintaining high relatedness within fruiting bodies, which may imply the existence of adaptations to reduce the risk of exploitation by maximizing the chance that cells aggregate with cooperative kin. There is evidence that *D. discoideum* actively segregates between kin and non-kin at least temporarily during aggregation [35, 36], though to what extent kin discrimination may protect against cheaters is unclear [37–39]. Nonetheless relatedness within fruiting bodies in nature is very high (Gilbert, Foster et al. 2007), probably owing to structure imposed by the way *D. discoideum* grows and disperses [40, 41]. Signatures of the frequency-dependent selection that often accompanies evolutionary conflict have been detected in genes known to affect cheating [42]. *D. discoideum* was also found to recognize and respond to the presence of non-kin within a chimeric aggregate with changes in gene expression, development, and dispersal behavior relative to clonal aggregates [43].

Despite multiple lines of evidence, however, some researchers have questioned the relevance of cheating to wild *D. discoideum.* As with any trait, directly proving that cheating is an adaptation is difficult, particularly when most studies of *D. discoideum* cheating involve laboratory-made chimeras. Non-fruiting obligate cheaters readily arise and prosper (at least over the short term) in experimental evolution experiments using *D. discoideum* under conditions imposing low relatedness [34], but zero were observed in a screen of 1039 spores isolated from 75 wild-collected fruiting bodies [24]. This may suggest that in nature, the benefits obligate cheaters reap by reducing stalk production in chimeras are too small or too infrequently realized to compensate for the disadvantages of being unable to produce a functioning stalk when there are no cooperative lineages to exploit – *D. discoideum* cheaters in nature are likely to be facultative cheaters. Results of genomic studies of *D. discoideum* genes differentially expressed in chimeras have been mixed – one study observed signatures of increased polymorphism and rapid evolution consistent with evolutionary conflict [44], while another did not [45]. Absent direct observation of cheating in nature, some researchers have proposed that apparent cheating is a laboratory artifact of little relevance to wild *D. discoideum*, and better explained by variation in non-social life history traits [46–48]. In this, cheating in *D. discoideum* is in good company with other well-supported social traits in microbes that have faced similar skepticism [49, 50].

In this study, we sought to infer whether or not *D. discoideum’s* apparent cheating was the result of adaptations to selective pressure to cheat (or resist cheating) in nature. We experimentally evolved multiple wild *D. discoideum* strains under conditions in which they never entered the social stage of their lifecycle and thus never had the opportunity to cheat. Without the opportunity to aggregate, selection on social traits like cheating should be relaxed, generally leading to losses in social function normally maintained by natural selection.

Key to this approach is the assumption that when a long-standing selective pressure is removed, past adaptations driven by that pressure are likely to be lost due to drift or, more likely, pleiotropic tradeoffs with other traits. Selection upon one trait will often indirectly impact other traits, and an organism’s traits often represent a compromise between mutually incompatible adaptations to different selective pressures. These compromises should tend to constrain the evolution of adaptations to overcome any particular selective pressure. An organism cannot, for example, evolve to be smaller to save energy and larger to avoid predation at the same time. If one selection pressure is relaxed, however, compromises are no longer necessary and constraints are lifted. For this reason, we should expect that relaxing a selective pressure should free an organism to lose traits that were adaptations to that pressure [51–54].

We can look for these sorts of losses and use them to infer selective pressures (and resultant adaptations) that we hypothesize are important but that are difficult to directly observe, such as the pressure that wild *D. discoideum* would experience if they were regularly exploited by cheaters. When *D. discoideum* is evolved in an environment where we know cheating is not relevant, it should lose adaptations related to cheating, but it cannot lose adaptations that it does not have. Therefore, loss of function when we make social conflict irrelevant in the lab is evidence that we have successfully relaxed a selective pressure that was relevant in nature. By contrast, if cheating in *D. discoideum* is the result of selective pressures on other, non-social traits and only *appears* to be cheating in an artificial laboratory environment, relaxation of selective pressures on the social stage should not affect it.

In this study, we experimentally evolve *D. discoideum* under conditions where cheating cannot occur, then measure the effects of experimental evolution on *D. discoideum’s* ability to cheat during the formation of chimeric fruiting bodies. In addition, we assayed the effects of experimental evolution on two other phenotypes to act as controls with which we could validate our logic. First, as an example of a trait that we were very confident experienced relaxed selection in our experiment, we assayed the distance travelled by *D. discoideum* slugs after aggregation but prior to fruiting body formation. The complexity of slug formation and migration and its ability to enable aggregates to move towards the soil surface and form fruiting bodies where they are in the best position to be dispersed is likely to be adaptive for *D. discoideum’s* in its natural environment [55]. In laboratory conditions where no slugs are formed, however, slug migration is irrelevant or even potentially maladaptive if there are tradeoffs between it and more useful traits. We expect slug migration distance to be uncorrelated, or negatively correlated with, fitness in the laboratory and so we expect to see our evolved *D. discoideum* lines evolve reduced migration distance relative to their ancestors.

As an example of a trait that we are confident would *not* experience relaxed selection in our experiment, we assayed the number of cells *D. discoideum* produced within 48 hours. Cell population is a product of traits – like growth rate and efficient use of resources – that we expect to be under strong selection both in nature and in the laboratory. In fact, the simplicity, mildness, and abundance of the laboratory environment is likely to result in *increased* selection to produce more cells more quickly relative to a natural environment full of abiotic and biotic threats and scarcity. We thus expect total cell production to be positively correlated with fitness in the laboratory, and do not expect to see loss of function in these traits – in the laboratory, our *D. discoideum* still use it, so they will not lose it. Together slug migration and cell production traits act as controls because they reflect a selective pressure that we experimentally relaxed and one that we did not relax, respectively. By comparing the effects of experimental evolution on cheating with its effects on these traits, we should be able to infer whether *D. discoideum* experiences selection pressure to cheat in nature.

## METHODS

### Culture conditions

We performed experimental evolution using SM/5 media [56] (2 g glucose (Fisher Scientific), 2 g BactoPeptone (Oxoid), 2 g yeast extract (Oxoid), 0.2 g MgCl_2_ (Fisher Scientific), 1.9 g KHPO_4_ (Sigma-Aldrich), 1 g K_2_HPO_5 (_Fisher Scientific), and for solid media 15 g agar (Fisher Scientific) per liter deionized water). To start a fresh culture of *D. discoideum*, we diluted spores from −80°C glycerol frozen stocks in KK2 buffer (2.25g KH_2_PO_4_ (Sigma-Aldrich) and 0.67g K_2_HPO_4_ (Fisher Scientific) per liter deionized water). We plated 1.0×10^5^ total spores onto an SM/5 plate along with 200µl *K. pneumoniae* food bacteria resuspended in KK2 to an OD_600_ of 1.5. To start a fresh culture of any bacterial strain, we streaked stocks from the minus 80°C freezer for isolation on SM/5 plates. We performed slug migration and spore production assays on nutrient-free starving agar plates.

### Antibiotic curing of *D. discoideum*

Many wild *D. discoideum* isolates are infected by *Paraburkholderia* symbionts which affect their fitness and behaviors [57]. To remove symbionts, 1.0×10^5^ *D. discoideum* spores of each clone were plated on SM/5 agar medium containing 30ug/mL tetracycline and 10ug/mL ciprofloxacin with 200µl of *K. pneumoniae* resuspended in KK2 to an OD_600_ of 1.5. We allowed plates to grow at room temperature under ambient light until formation of fruiting bodies (3-5 days). We collected spores as above, then diluted and plated again as above. We then collected spores and performed spot test assays (described in [58]) and PCR using *Paraburkholderia-*specific primers [59] to verify successful curing.

### Experimental evolution

Three replicate lines each of ten strains (**Supp. Table 1)** were plated on SM/5 plates. We incubated all lines at room temperature under ambient light and transferred 0.5% of the population to fresh plates every 48 hours. We performed transfers by first harvesting all cells into 10mL KK2 buffer using gentle pipetting and scraping of the agar surface. We then thoroughly vortexed the resulting suspensions, diluted them 200-fold, and plated 100µL onto fresh plates with 200µL of an OD_600_ = 1.5 *K. pneumoniae* suspension to serve as food. The 48-hour transfer interval was selected to preempt *D. discoideum’s* fruiting stage and prevent direct selection on social traits. Every fifth transfer, we additionally froze 1mL of the undiluted suspension of harvested cells at −80°C with 60% glycerol.

Following experimental evolution over 30 transfers, *D. discoideum* lines were checked for cross-contamination using fragment analysis. We extracted DNA from 100µL of the undiluted suspension of harvested cells using CHELEX resin beads and amplified using fluorescently-tagged PCR primers specific to highly variable microsatellite loci known to differ in length between *D. discoideum* strains [60]. Fragment analysis of resulting amplicons was performed by Genewiz and evolved strains were compared to ancestors. No cross-contamination was detected.

### Cheating assays

To assay cheating, we determined the proportion of fluorescent spores in fruiting bodies developing from an initial 50:50 mix of a strain of interest and the RFP-labelled control strain RFP-NC28.1. We assayed cheating in chimeras comprising lines of interest and a labelled control strain to reflect the natural context where a cheater must compete against an unrelated strain. We plated 2×10^5^ spores for each strain onto SM/5 agar plates. During mid-log stage (approximately 34-36 hours), we collected vegetative cells and washed three times with cold KK2 buffer, counted using a hemacytometer, and diluted each suspension to 10^8^ cells/mL. We combined equal volumes of the focal strain and the labelled control strain and gently mixed to get 50:50 mix suspensions. We prepared UV-sterilized 13mm^2^ AABP 04700 (Millipore) filter squares. We pipetted 15µL of the 50:50 mix suspension into the center of each of 3 filter squares pre-dampened with KK2. We transferred filters onto KK2 (non-nutrient) agar plates to initiate immediate aggregation and development and incubated plates for 5 days at room temperature.

After incubation, we examined and selected 2 filters from each plate to assay. We prioritized filters that showed no evidence of slugs having escaped onto the surrounding agar – otherwise selection was random. We collected each filter with the fruiting bodies that had grown atop them with sterile forceps into 500µL KK2 buffer, vortexed thoroughly, and took photographs with bright-field and fluorescence microscopy. We captured at least 5 fields for each sample (representing around 1000-2000 spores). Each assay was performed on three separate days.

We counted total spores and fluorescent spores using Fiji [61]. We first manually converted micrographs into binary images using a brightness threshold set individually for each image to account for minor differences in contrast and brightness between samples. We then used Fiji’s Count Particles function, filtering for particles between 15-200 um^2^ and between 0.5 and 1.0 circularity (settings which consistently resulted in very similar results to manual counting).

We determined proportion of fluorescent spores by dividing the number of fluorescent spores (count from fluorescent image) by the total spores (count from brightfield image). We also determined a percentage fluorescence for the RFP-NC28.1 control for each assay and divided this proportion out of the results for the 50:50 mix samples in order to compensate for incomplete labelling.

### Slug migration assays

In order to compare how experimental evolution affected slug migration distance, we performed assays on ancestral and evolved lines. Each assay was performed in triplicate on non-nutrient agar plates (13 cm diameter). We marked a 10 cm secant line on the back of each plate. We plated 50μL of an OD_600_=50.0 suspension of *K. pneumoniae* in KK2 buffer containing 10^7^ *D. discoideum* spores along the secant line. We allowed the loaded sample to dry and wrapped the plates individually in aluminum foil. On each wrapped plate, we made a small pinhole opposite the starting line through which light could enter. We placed the wrapped plates on the laboratory bench under a light source and left them undisturbed for 8 days. At the end of the 8 days, we unwrapped the plates and photographed using a Canon EOS 5D Mark III camera.

We used Fiji [61] to perform image processing and obtain slug migration distances. First, we scaled images and overlayed a 1cm x 1cm grid. We marked fruiting bodies on each image and measured their distance from the starting line.

### Cell count assays

To assay cell populations, we performed assays on ancestral and evolved lines. For each ancestral and evolved line, we plated 1×10^5^ spores on a fresh SM/5 nutrient agar plate, along with 200µl of an OD_600_=1.5 suspension of *K. pneumoniae* food bacteria. Plates were incubated at room temperature. After 48 hours, all cells were collected into a suspension of KK2 buffer by pipetting buffer onto each plate’s surface and gently agitating with a pipette tip. The resulting suspensions were then vortexed and counted using a hemocytometer to determine total cell populations on each plate. Conditions, timing, and process of cell count assays were identical to those used during the experimental evolution process. Each assay was performed in triplicate.

### Spore count and sporulation efficiency assays

We assayed spore production using the plates prepared for the slug migration assays. One day after plates from the slug migration assays were imaged, we collected the fruiting bodies from each plate into KK2 buffer. We counted the spores within the resulting suspensions under a light microscope using a hemocytometer.

To test the possibility that apparent reductions in cheating were the result of differences in the efficiency of sporulation of evolved *D. discoideum* lines, we also performed spore production assays on ancestral and evolved lines on plates without food bacteria. For each ancestral and evolved line, we plated 1.5×10^6^ amoebae onto UV-sterilized 13mm^2^ AABP 04700 (Millipore) filter squares on a fresh non-nutrient KK2 plate without exogenous food. Under these conditions, amoebae immediately starve and enter the fruiting stage. After 96 hours, we collected each filter into a tube with 500µl KK2 buffer, vortexed, and counted using a hemocytometer to determine the total spore production of each plate. By immediately inducing the fruiting stage and leaving no opportunity for growth, this assay effectively isolates differences in the efficiency with which *D. discoideum* lines could convert amoebae into spores.

### Statistical analysis

We performed analyses using R version 4.0.4 (R Core Team, 2015) with the *lme4* package (Bates et al., 2015) and the *emmeans* package (Lenth, 2022).

To analyze changes in spore production and slug migration distance, we used linear mixed effects models with treatment (ancestor vs evolved) as a fixed effect and strain as a random effect. The model used to analyze slug migration was Average Migration Distance ∼ Treatment + (1|Strain/Line). In the slug migration assays, in order to account for fruiting bodies that developed at the starting line (migration distance = 0), we weighted each plate’s contribution to the model by the fraction of spores from fruiting bodies with nonzero migration distances. This had the effect of reducing the average migration distance on each plate appropriately by the fraction of spores that did not migrate beyond the starting zone. The model used to analyze cell count and spore count data was Total Counted ∼ Treatment + (1|Strain/Line).

To analyze changes in representation among spores in chimeric fruiting bodies, we used a two-tailed, two-proportion Z test.

In order to convey the most biologically meaningful insight, we have reported our results in terms of 95% compatibility intervals in addition to p-values [62].

## RESULTS

### Cheating assays

For each *D. discoideum* line, we combined amoebae of the line of interest with amoebae of the control strain RFP-NC28.1 at a 50:50 ratio, then measured the proportion of spores within the resulting chimeric fruiting bodies belonging to the control strain. Ancestral *D. discoideum* strains made up an average of 54.27% of spores in chimeric fruiting bodies, suggesting that overall the wild strains could cheat RFP-NC28.1. In chimeras made from evolved *D. discoideum* strains and RFP-NC28.1, evolved *D. discoideum* strains were significantly less well represented than their ancestors had been (χ^2^(1)=175.34, p<0.001), contributing an average of 51.07% of spores. Possible values for the true decrease most compatible with our data ranged from −2.65% to −3.56% (95% CI) **(Figure 2)**.

**FIGURE 2.**
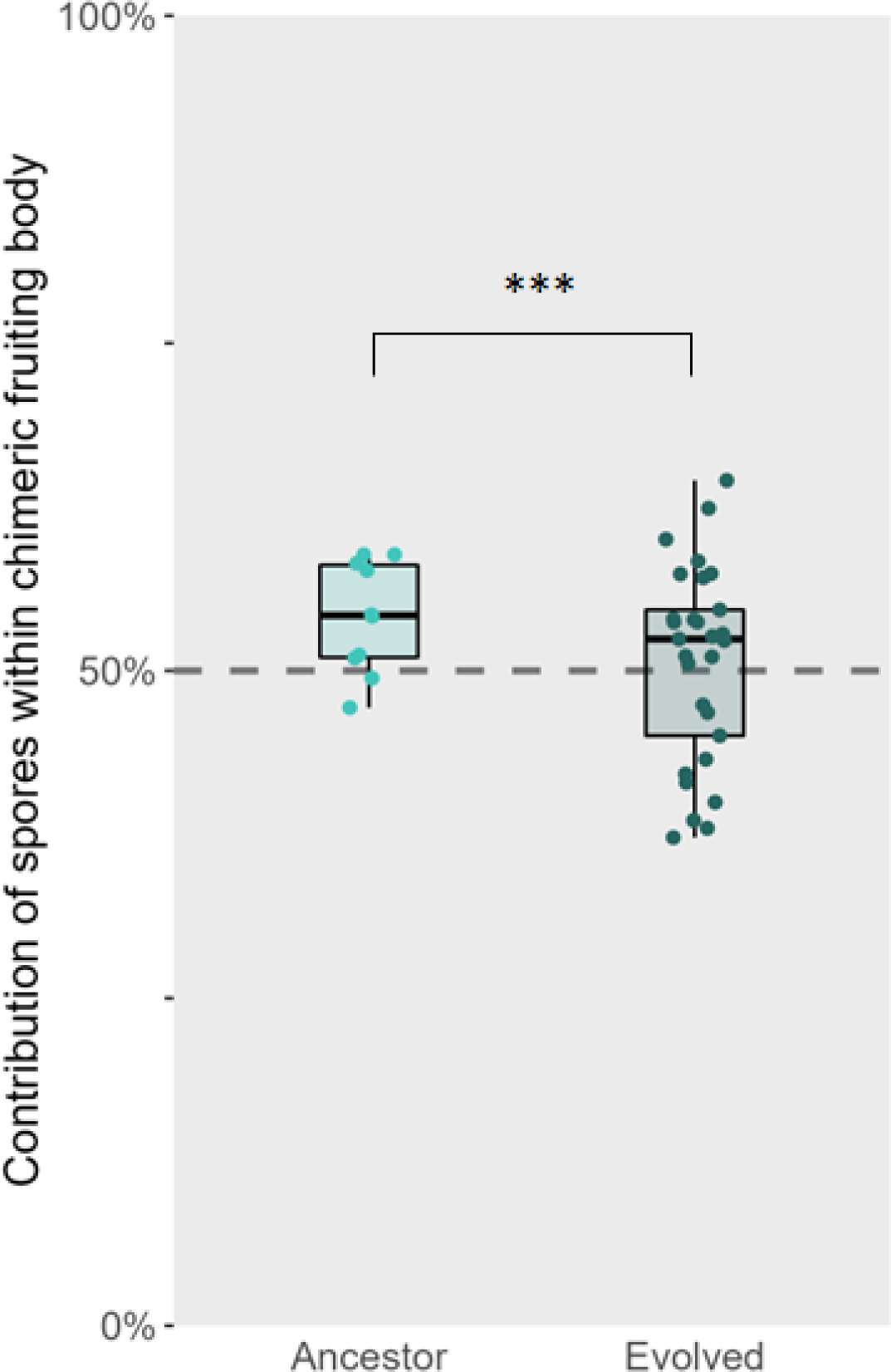
Experimentally evolved *D. discoideum* evolve reduced ability to cheat. Spores of ancestral *D. discoideum* strains are slightly overrepresented within chimeric fruiting bodies made by combining equal numbers of the focal strain and a labelled control strain. When competed against the same control strain, evolved *D. discoideum* strains contribute a lower percentage of spores within fruiting bodies compared to their ancestors, suggesting that experimental evolution resulted in reduced cheating ability (or reduced ability to resist being cheated upon).

The reduced representation of evolved lines among spores within chimeras was not the result of evolved lines being less capable of producing spores than their ancestors. In growth assays, we found that when inoculated clonally on SM/5 plates and provided exogenous food, ancestral *D. discoideum* produced an average of 6.9×10^6^ spores on a plate over the course of 8 days. Experimentally evolved *D. discoideum* lines produced an average of 1.5×10^6^ (+21.9%) *more* total spores than their ancestors (t(28.99)=2.392, p=0.0235). Possible values for the true increase most compatible with our data ranged from 2.5×10^5^ to 2.7×10^6^ (+3.7% to +40.1%) (95% CI) (**Supp. Fig. 1).** We also did not detect a significant difference in sporulation efficiency between amoebae from ancestral and evolved *D. discoideum* lines inoculated clonally on SM/5 plates *without* exogenous food (t(28.09)=0.421, p=0.677). Under these conditions, which should eliminate any effect of differences in vegetative cell fitness between lines, ancestral *D. discoideum* produced an average of 9.31×10^5^ spores per plate after 4 days. Experimentally evolved *D. discoideum* produced an average of 9.57×10^5^ (+2.7%) spores per plate over the same timeframe. Possible values for the true change most compatible with our data ranged from 8.38×10^5^ to 1.08×10^6^ (−10.1% to +15.9%) (95% CI) **(Supp. Fig. 2).**

### Slug mobility assays

We measured the effects of experimental evolution on slug migration by assaying the average distance travelled by slugs produced by ancestral and evolved lines from a starting position. Ancestral *D. discoideum* slugs migrated an average of 4.01cm from the starting line before fruiting. Experimentally evolved *D. discoideum* slugs migrated less far, moving an average of 3.52cm (−12.2%) less far than their ancestors (t(109)=-2.969, p=0.00367). Possible values for the true reduction in migration distance most compatible with our data ranged from 0.82cm to 0.17cm (−20.4% to −4.2%) (95% CI) **(Figure 3B)**.

**FIGURE 3.**
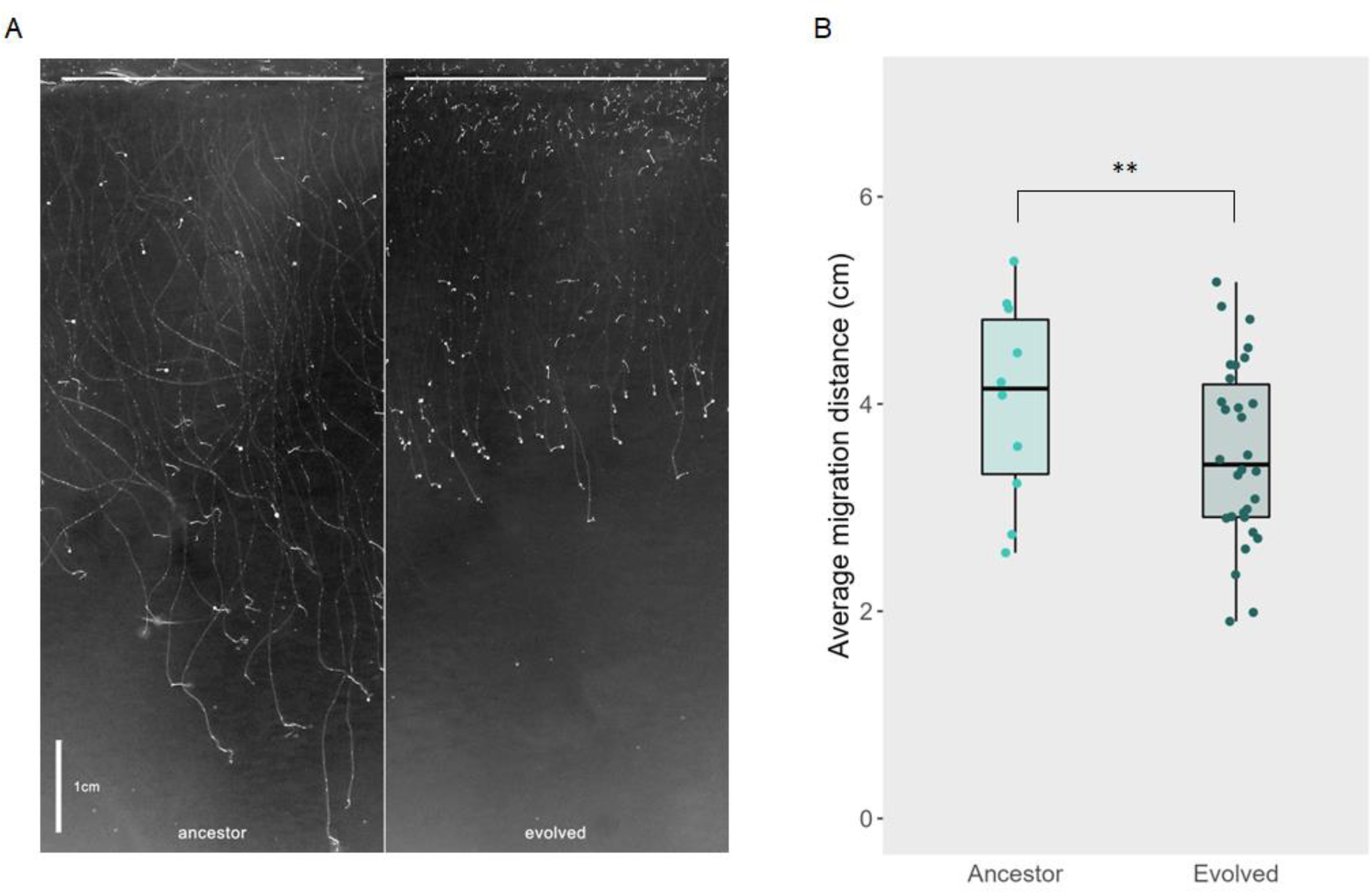
Experimentally evolved *D. discoideum* evolve reduced slug mobility. **A)** Representative image of slug mobility assays. *D. discoideum* suspensions were plated along the starting line (top of the figure). Slugs migrated towards a light source (bottom of figure). B) *D. discoideum* strains evolved without the opportunity to aggregate evolved to migrate shorter distances during the slug stage than their ancestors.

### Cell count assays

We measured the effects of experimental evolution on *D. discoideum’s* ability to reproduce in the laboratory by comparing the total cell populations reached by ancestor and evolved lines in 48 hours. Ancestral *D. discoideum* reached an average population of 1.9×10^8^ cells per plate after 48 hours. Experimentally evolved *D. discoideum* produced an average of 5.0×10^7^ (+26.3%) more cells than their ancestors within the same timeframe (t(2.9)=2.366, p=0.0249). Possible values for the true increase most compatible with our data ranged from 8.5×10^6^ to 9.1×10^7^ (+4.5% to +47.9%) (95% CI) **(Figure 4).**

**FIGURE 4.**
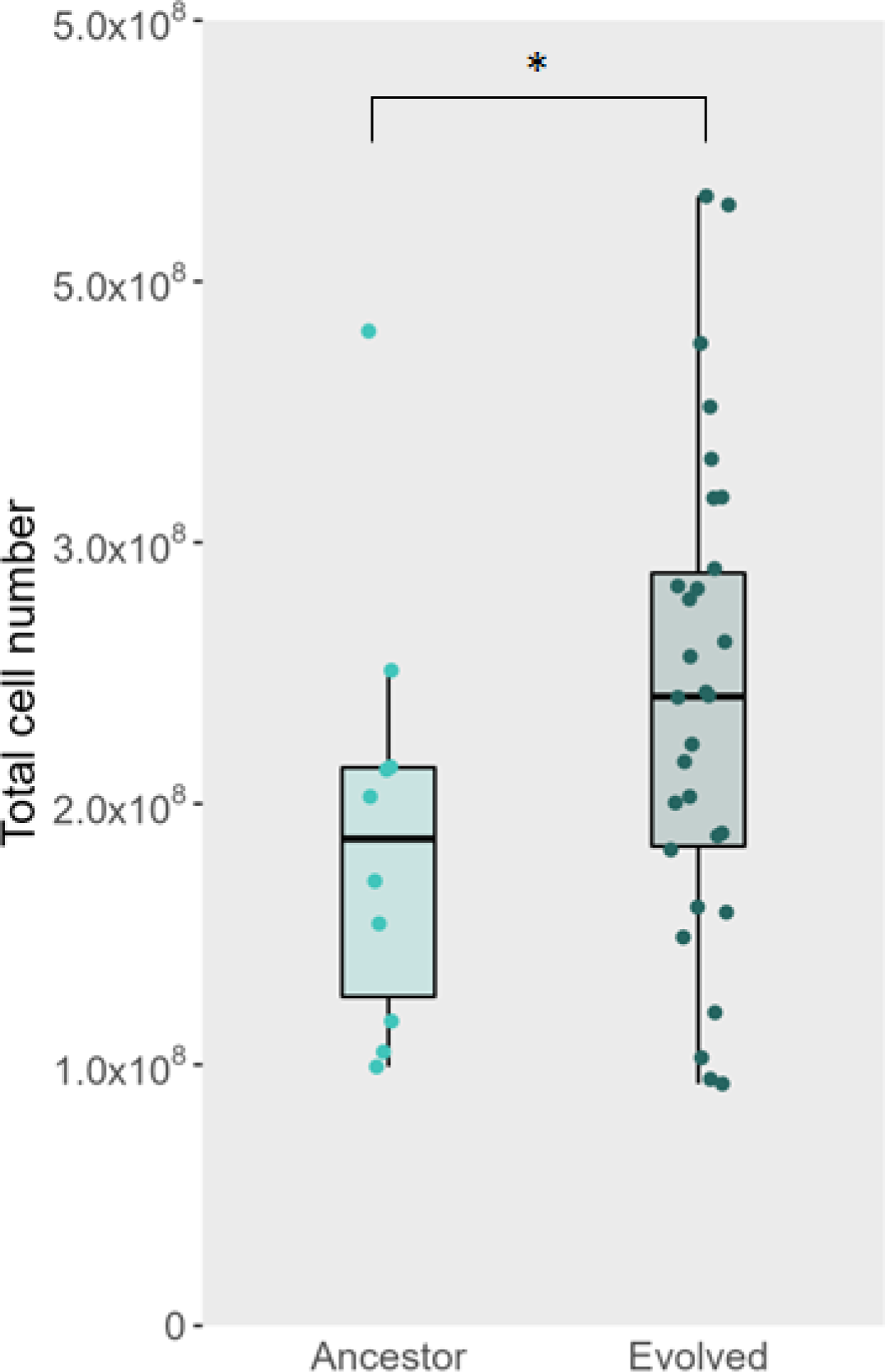
Experimentally evolved *D. discoideum* strains produce more cells. **A**) *D. discoideum* strains evolved without the opportunity to aggregate produce significantly more cells than their ancestors within a 48 hour period.

## DISCUSSION

*D. discoideum* is a useful model organism for studying cooperation and conflict. Its social cycle requires potentially unrelated cells to cooperate, but also creates incentive for cells to exploit one another. While multiple lines of evidence attest to some *D. discoideum* strains’ ability to exploit one another in the laboratory, interpreting the relevance of cheating to *D. discoideum* in nature is not trivial. In this respect, cheating in *D. discoideum* echoes other prominent examples of microbial sociality [49, 50] and illustrates a general challenge in studying adaptations in any organism. It is easier to prove that an organism has a particular trait than it is to be certain what selective pressures (if any) drove the trait’s evolution, and doubly so if the organism is too small to directly observe in its natural habitat.

This study attempts to shed light on the adaptive value of cheating in *D. discoideum* in nature by experimentally evolving wild strains of *D. discoideum* under laboratory conditions in which the social cycle – and thus cheating – is prevented. This experiment hinges on the idea that relaxing selective pressures an organism faces in nature will tend to result in loss of adaptations maintained by those pressures [53, 66–69]. We looked for changes in experimentally evolved *D. discoideum’s* ability to cheat that would suggest it had experienced selective pressure to cheat (or resist cheating) in its natural environment. In addition, we assayed slug migration distance and cell number, two phenotypes for which we had opposite *a priori* predictions.

We found that compared to their ancestors, experimentally evolved *D. discoideum* were less capable cheaters within chimeric fruiting bodies (**Figure 2)**. This result is what we would expect if our ancestral *D. discoideum* strains had adaptations to make them better cheaters (or more resistant to being cheated upon) that atrophied when we relaxed the selective pressures that were maintaining them. However, an alternative explanation for the poorer representation of evolved lines within chimeras is that evolved lines might have simply become less capable of producing spores due to drift or pleiotropy during evolution in an environment where spore production did not occur. When mixed in chimeras with a wild type strain, these less productive strains could appear to undercontribute to spore production even with no cheating involved. To test this possibility, we assayed spore production of clonal populations of ancestral and evolved lines and found that evolved lines produced *more* spores on average, rather than fewer **(Supp. Figure 1).** To account for the potential confounding effect of increased vegetative cell fitness on spore production, we also assayed sporulation efficiency directly and observed no difference between ancestral and evolved lines **(Supp. Figure 2).** Ultimately, our evolved lines are at least as good at producing spores as their ancestors, and often better. Since this would tend to make them appear to be *better* cheaters in our cheating assays, we suspect that the reduction in cheating we observed is, if anything, an underestimate, and that the best explanation is that we were observing losses of adaptations no longer being maintained by selection.

This interpretation is further strengthened by our results for slug migration and cell number, which act positive and negative controls, respectively, for our method. Slugs produced by experimentally evolved *D. discoideum* lines migrated less far than those produced by their ancestors **(Figure 3).** As slug migration is certainly an adaptive trait in nature, we expected our ancestral lines to have adaptations that enhanced it. Accordingly, when we relaxed selective pressure on these adaptations by preventing slug formation entirely, we expected to see evolved lines form less functional slugs for the same reason we are suggesting that they became less functional cheaters. By contrast, we found that evolved *D. discoideum* produced more cells than their ancestors (**Figure 4**), likely as the result of selection for higher growth rates and/or more efficient use of resources in the laboratory during the vegetative stage. This reflects that the laboratory environment – where food is plentiful and conditions are mild – behooves cells to invest in outcompeting neighbors rather than protect themselves from no-longer-relevant hazards that they would face in nature. Cell number remained at least as relevant during our experimental evolution experiment as it would have been in nature, and thus we would not expect the reduction in this trait that we observed in the cheating and slug migration distance traits.

Adaptations are not free, and adapting to enhance one trait will often come at the expense of unrelated traits that may be adaptive for other reasons [63–65]. While positive pleiotropic interactions exist between some traits, it is more common for unrelated functions to trade off negatively for the same reason that detrimental mutations are more common than beneficial mutations – there are more ways to break a system than there are to improve it [52]. The prevalence of antagonistic pleiotropy between traits means that adaptations often represent some balance between competing constraints. If *D. discoideum* has adaptations that enhance its ability to cheat (or resist being cheated), they are likely to come at some cost to adaptations to other selective pressures. When *D. discoideum* is experimentally evolved under conditions where it is no longer selected to cheat, these tradeoffs should incentivize it to lose cheating adaptations. By contrast, if cheating is not selected for in nature, *D. discoideum* should not have adaptations supporting it, and moving cells to an environment where cheating continues not to be adaptive should not change anything.

The results of our cheating assays suggest that wild *D. discoideum* does have adaptations related to social conflict which they lost when evolved in an artificial environment in which cheating was necessarily irrelevant to their fitness. The removal of any selective pressure on the social stage freed experimentally evolved lines to evolve based upon non-social selective pressures alone. Our results are consistent with past molecular evolution studies implying *D. discoideum* genes affecting cheating have an evolutionary history driven by social conflict [42, 44]. *D. discoideum* likely does cheat in nature.

Microbes lead complicated lives obscured from us by alien tininess. Understanding even some apparently central aspects of their biology is a complex task demanding multiple approaches and careful interpretation. *D. discoideum’s* social cycle has been the subject of interest for decades, and yet how exactly it fits into this microbe’s life in nature continues to inspire debate. The results of this study support the idea that some wild *D. discoideum* strains experience enough social conflict to have evolved adaptations to it. Further, the approach we have employed here – looking for otherwise inscrutable adaptations by evolving them away in lab – should be applicable to a wide variety of traits in a wide variety of organisms. This approach can usefully supplement other approaches in researchers’ pursuit of a more complete understanding of adaptation.

## ACKNOWLEDGEMENTS

We thank members of the Queller/Strassmann lab, Jonathan Losos, Carlos Botero, and Fred Inglis for helpful discussion during the writing of this paper. This material is based upon work supported by the National Science Foundation under Grants IOS 1656756 and DEB 1753743, and DEB 2237266.

---

**Supplemental Table 1.**

| Strain | Alt ID | Species | Origin | Date collected | Note |
| --- | --- | --- | --- | --- | --- |
| QS6 | V77B | Dictyostelium discoideum | Mountain Lake Biological Station, VA, USA | 9/25/2000 |  |
| QS9 | V330D2 | Dictyostelium discoideum | Mountain Lake Biological Station, VA, USA | 10/15/2000 |  |
| QS11 | V324B1 | Dictyostelium discoideum | Mountain Lake Biological Station, VA, USA | 7/21/2004 |  |
| QS18 | V342B2 | Dictyostelium discoideum | Mountain Lake Biological Station, VA, USA | 10/15/2000 |  |
| QS69 | DD10C2 | Dictyostelium discoideum | Houston Arboretum, TX, USA | 5/18/2004 |  |
| QS70 | HD25A1 | Dictyostelium discoideum | Houston Arboretum, TX, USA | 7/15/2004 |  |
| QS159 | 14/7 | Dictyostelium discoideum | Mountain Lake Biological Station, VA, USA | 1/5/2008 |  |
| QS161 | UB1 | Dictyostelium discoideum | Mountain Lake Biological Station, VA, USA | 1/5/2008 |  |
| QS395 | V319B1 | Dictyostelium discoideum | Mountain Lake Biological Station, VA, USA | 10/15/2000 |  |
| QS859 | 7P7.1 | Dictyostelium discoideum | Mountain Lake Biological Station, VA, USA | 7/23/2014 |  |
| QS48-RFP | NC28.1 | Dictyostelium discoideum | Linville Falls, NC, USA | 7/3/2001 | RFP-labelled control strain used in cheating assays |

**SUPPLEMENTARY FIGURE 1.**
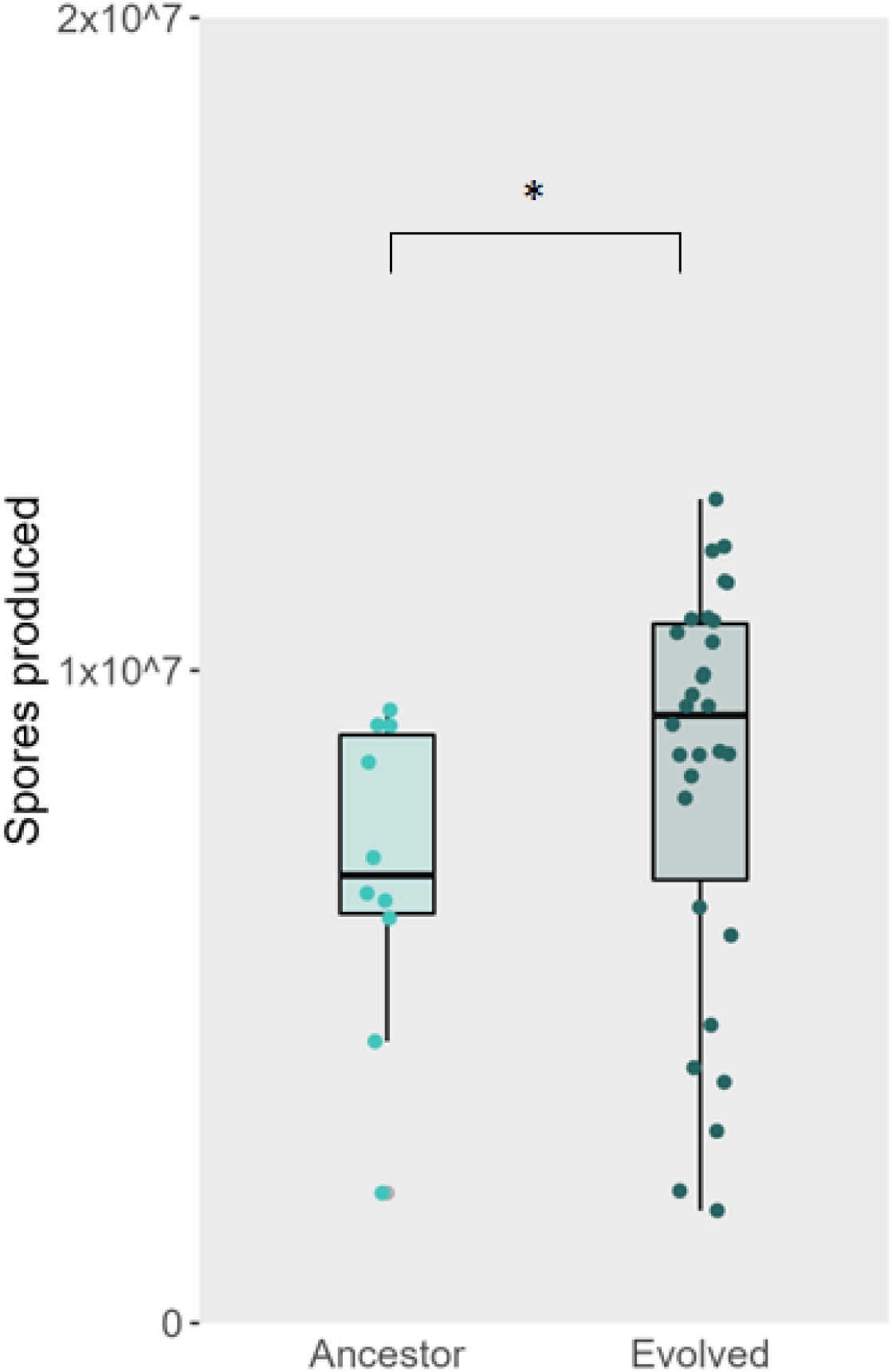
Experimentally evolved *D. discoideum* strains evolve increased spore production. *D. discoideum* strains evolved without the opportunity to aggregate produce significantly more spores than their ancestors. This difference may reflect experimentally evolved lines reaching higher populations during the vegetative stage prior to fruiting (see main paper figure 4).

**SUPPLEMENTARY FIGURE 2.**
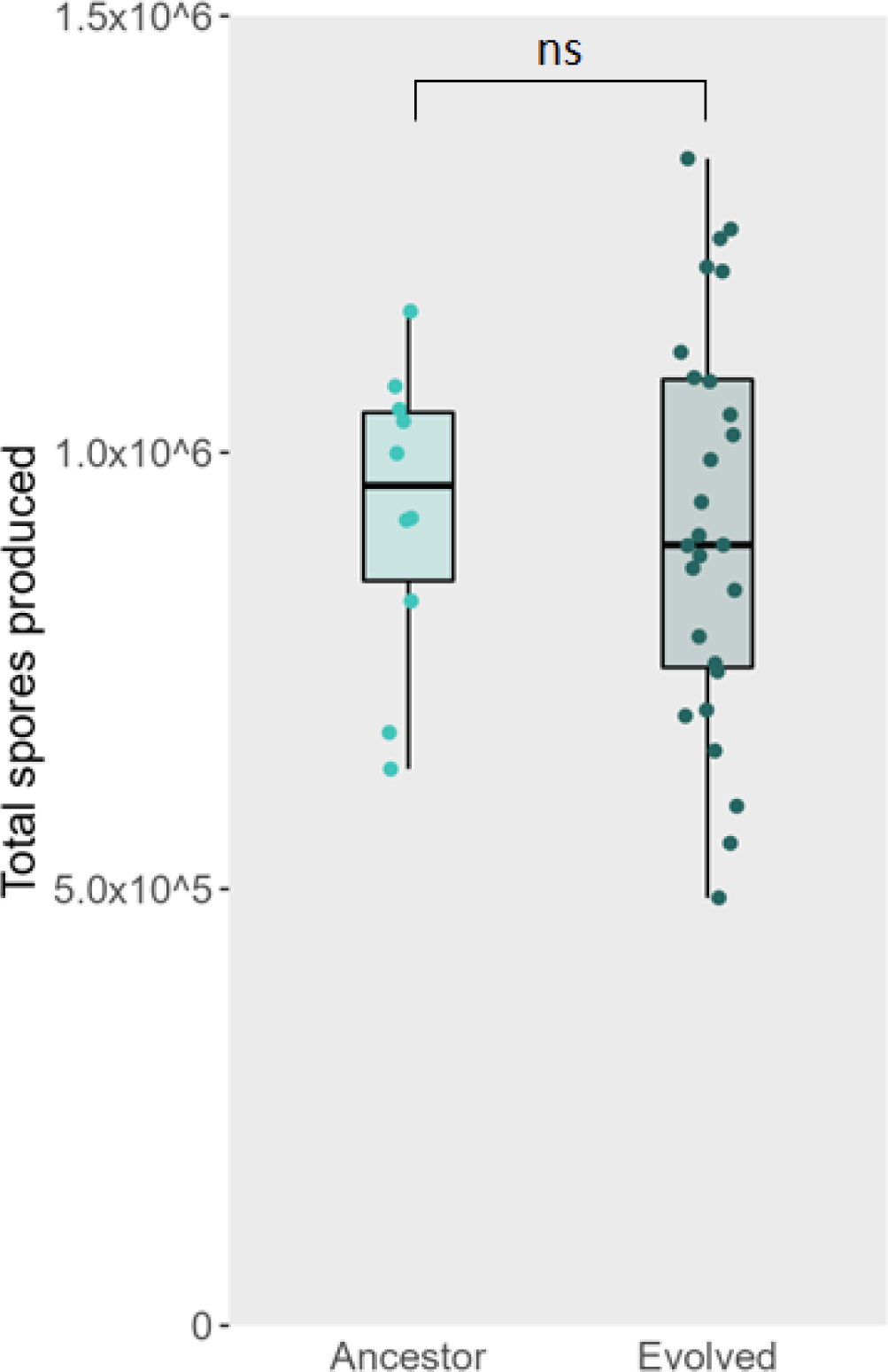
Experimentally evolved *D. discoideum* lines are not less efficient at sporulating. Amoebae were plated on filters without exogenous food and allowed to fruit. Mature fruiting bodies were then collected into KK2 suspensions and spores were quantified. This assay eliminates any impact of differences in lines’ fitness during the vegetative stage, and therefore act as a measure of the efficiency by which amoebae can be converted into spores. Experimentally evolved *D. discoideum* lines did not differ significantly in sporulation efficiency compared to their ancestors.

